# Brain oscillatory processes related to sequence memory in healthy older adults

**DOI:** 10.1101/2023.09.28.559534

**Authors:** Nina M. Ehrhardt, Agnes Flöel, Shu-Chen Li, Guglielmo Lucchese, Daria Antonenko

## Abstract

Sequence memory is subject to age-related decline, but the underlying processes are not yet fully understood. We analyzed electroencephalography (EEG) in 21 healthy older (60-80 years) and 26 young participants (20-30 years) and compared time-frequency spectra and theta-gamma phase-amplitude-coupling (PAC) during encoding of the order of visually presented items. In older adults, desynchronization in theta (4-8 Hz) and synchronization in gamma (30-45 Hz) power did not distinguish between subsequently correctly and incorrectly remembered trials, while there was a subsequent memory effect for young adults. Theta-gamma PAC was modulated by item position within a sequence for older but not young adults. Specifically, position within a sequence was coded by higher gamma amplitude for successive theta phases for later correctly remembered trials. Thus, deficient differentiation in theta desynchronization and gamma oscillations during sequence encoding in older adults may reflect neurophysiological correlates of age-related memory decline. Furthermore, our results indicate that sequences are coded by theta-gamma PAC in older adults, but that this mechanism might lose precision in aging.

## 1. Introduction

In addition to a general decline of memory functions, the ability to remember the order of events, i.e., sequence memory, is impaired in aging (Li et al., 2010; Newman et al., 2001; Rotblatt et al., 2015; Seewald et al., 2018). Brain oscillatory processes during encoding and binding of memory associations contributing to the decline are not yet fully understood.

In young adults, formation of episodic memories has been associated with localized synchronous neuronal activity as indexed by power in different frequency bands, as well as inter-frequency coupling (for reviews see e.g., Hanslmayr et al., 2016; Köster and Gruber, 2022; Nyhus and Curran, 2010). Power decreases were observed in the frequency range of 10-30 Hz (alpha/beta band) during encoding, interpreted as desynchronization of local groups of neurons (Buzsaki and Draguhn, 2004) and associated with successful retrieval (Fellner et al., 2013; Griffiths et al., 2021; Karlsson et al., 2020; Martín-Buro et al., 2020). On the other hand, in higher frequencies (above 30 Hz, i.e. the gamma band) increases in power have been associated with successful encoding (Fellner et al., 2019; Friese et al., 2013; Osipova et al., 2006; Roehri et al., 2022; Sederberg et al., 2006). In addition, the temporal coordination between theta phase and gamma amplitude, i.e. phase-amplitude coupling (PAC), has been linked to successful episodic memory performance (Friese et al., 2013; Griffiths et al., 2021; Heusser et al., 2016), providing a mechanism to represent the serial order of encoded stimuli (Heusser et al., 2016).

These findings regarding oscillatory processes during encoding in young adults - from local power changes in different frequency bands to interactions between oscillations of different frequencies - have been integrated in the Sync/DeSync-model (Hanslmayr et al., 2016; Parish et al., 2018). In this model, the desynchronization in lower frequencies (< 30 Hz) leads to disinhibition of cortical regions, thereby allowing the perceptual representation of to-be-remembered stimuli (Hanslmayr et al., 2016; Hanslmayr et al., 2012; Parish et al., 2018). The to-be-remembered items are then thought to be represented by gamma spikes in the cortex as input to the hippocampus for forming memory associations (Griffiths et al., 2021; Lisman and Jensen, 2013; Nyhus and Curran, 2010).

In older adults, such a comprehensive view integrating multiple findings of interacting oscillations during the formation of episodic memories is still missing. While some studies investigated the association of frequency bands or theta-gamma PAC with episodic (Crespo-Garcia et al., 2012; Després et al., 2017; Karlsson et al., 2022; Karlsson and Sander, 2023; Karlsson et al., 2020; Lithfous et al., 2015; Sander et al., 2020; Strunk and Duarte, 2019; Winterling et al., 2019) or working memory (Reinhart and Nguyen, 2019), the aging related effects have not been investigated specifically with respect to sequence memory.

To investigate the interplay of oscillations at different frequencies during the formation of sequence memories, we analyzed electroencephalography (EEG) from 21 older (age range: 60-80 years) and 26 young adults (age range: 20-30 years) during an adapted version of a sequence memory task investigated by Heusser et al. (2016). First, we aimed to compare time-frequency spectra in common frequency bands (theta, alpha/beta, gamma) during early (perceptual) and late (binding) phases of encoding for later correctly and incorrectly remembered sequences between young and older adults. We hypothesized that the differential synchronization and desynchronization processes, postulated for young adults in the Sync/Desync model (Hanslmayr et al., 2016; Parish et al., 2018), would be disturbed in older adults such that there will be less desynchronization in lower frequency bands (theta, alpha/beta) as an early response to a stimulus compared to young adults. Additionally, the following synchronization in gamma frequency in a later time window will be lower in older compared to young adults. Next, we examined the coding of sequences through coupling of gamma amplitude to theta phase for young and older adults. We hypothesized that the pattern of coupling between gamma amplitude and theta phase that was found for young adults (Heusser et al., 2016) would be disturbed in older adults, indicated by a less consistent modulation of gamma power by theta phase according to the picture position within a sequence.

## 2. Methods

### 2.1 Participants

Thirty-two healthy older (21 woman; average age = 68 years, range = 61-79 years) and thirty-two healthy young (19 woman; average age = 24 years, range = 20-30 years) adults participated in this study. Participants were right-handed, fluent in German, without history of neurological or psychiatric disease (self-reported and/or Geriatric Depression Score ≥ 6 (Yesavage et al., 1982) for older adults) and had normal or corrected-to-normal hearing and vision. After EEG preprocessing (see below), data from 21 older and 26 young adults was included in the analyses. Besides age the groups did not differ significantly from each other regarding the demographics (see ***Table 1***).

**Table 1.** Demographic information of the participants.

|  | Older adults<br>(n = 21) | Young adults<br>(n = 26) |  |
| --- | --- | --- | --- |
|  | Mean (SD) |  |  |
| Age in years | 67.76 (5.97) | 24.27 (2.41) | $t(45) = 33.93, p < .001$ |
| Gender (n, % female) | 15 (71.43) | 14 (53.85) | $\chi^2 = 0.87, p = 0.35$ |
| Education in years | 15.36 (2.13) | 16.35 (1.78) | $t(45) = -1.73, p = 0.09$ |
| EHI | 92.38 (12.21) | 94.42 (16.75) | $t(45) = -0.47, p = 0.64$ |

*Note.* SD = standard deviation. EHI = Edinburgh Handedness Inventory.

Older participants with impaired cognition (subjectively reported in screening interview or identified in baseline assessment as z-scores < -1.5 in any of the tests in the neuropsychological test battery of the Consortium to Establish a Registry for Alzheimer’s Disease, CERAD, https://www.memoryclinic.ch) were not included (see ***Supplementary Table*** for descriptive neuropsychological information of older participants). Participants performing at chance level (<60% correct; n=14) in the memory paradigm (see below) were also not included to ensure compliance with task instructions (Filmer et al., 2017; Hsieh et al., 2011; Violante et al., 2017), except three older participants who reached <60% correct overall due to low performance only in the last two blocks (likely reflecting exhaustion rather than non-compliance). All participants provided written informed consent and received reimbursement for their participation. The study was conducted in accordance with the declaration of Helsinki, and approved by the ethics committee of University Medicine Greifswald. Data from the older adults has been used to identify stimulation parameters for a study investigating the effects of transcranial alternating current stimulation (tACS) on episodic memory in healthy older adults (Ehrhardt et al., 2023).

### 2.2 Experimental Paradigm

Participants performed an adapted version of a sequence memory task (Heusser et al., 2016) (see ***Figure 1***).

**Figure 1.**
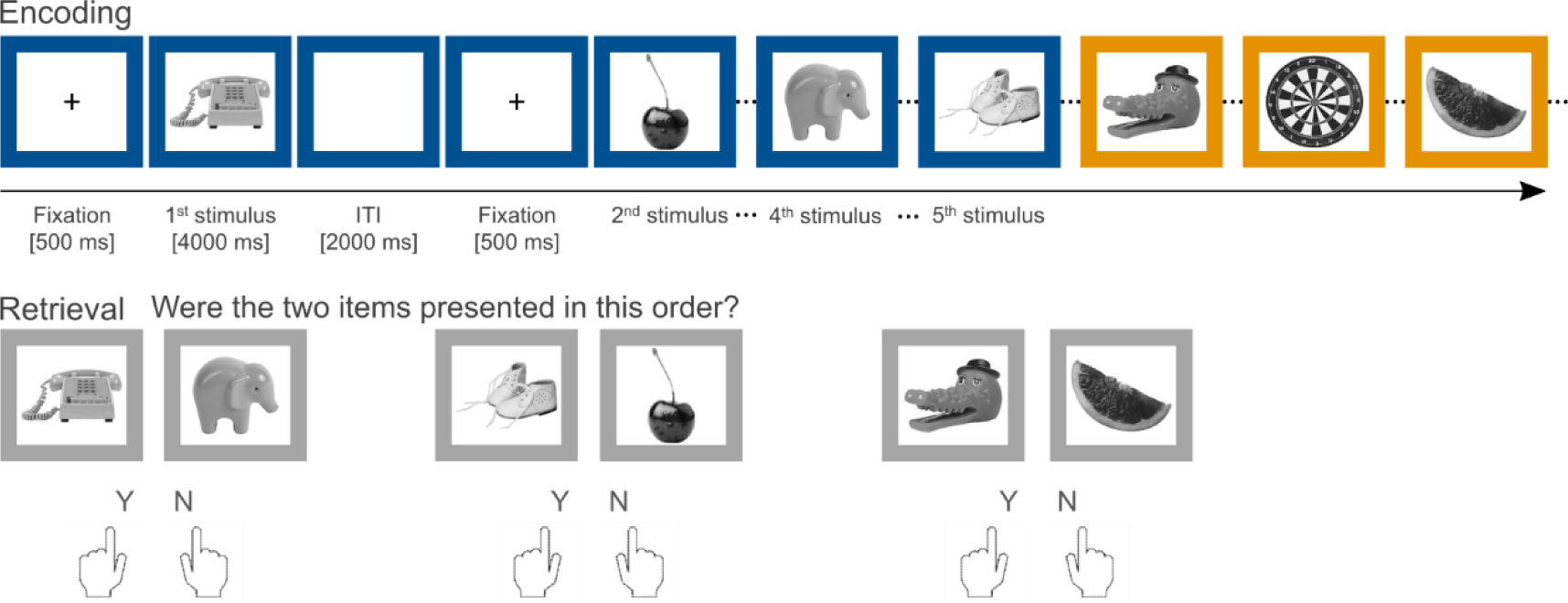
The sequence memory task. During encoding, participants saw grey-scale pictures presented sequentially in colored frames. Each picture was on screen for 4000 ms preceded by a fixation cross at the center of the screen for 500 ms and followed by an inter-trial interval of 2000 ms. After a list of five pictures, the color of the frame changed. During retrieval after the presentation of four lists, two of the previously shown pictures were presented next to each other in a grey frame and participants indicated whether the pictures were presented in the right temporal order by button press. In total, 12 blocks of picture lists and corresponding retrieval phases were presented.

Grey-scale pictures of non-human objects without negative valence (e.g., weapons, spiders) and not containing any text (retrieved from http://cvcl.mit.edu/MM/uniqueObjects.html) were presented sequentially in colored frames. Participants were asked to remember the temporal order of the presented pictures. In total, 12 blocks of pictures were presented, each followed by a retrieval phase. Every encoding trial consisted of a fixation cross at the center of the screen for 500 ms, a picture displayed for 4000 ms, and an inter-trial interval of 2000 ms. For deeper encoding participants were asked to apply two strategies. First, imagining the shown object in the color of the frame and indicating whether the color fits the picture by button press. Second, imagining the object interacting with the previous object. A block consisted of four lists with five pictures, the color of the frame changing with every list. Each block was followed by the retrieval phase in which sequence memory was tested. In the retrieval phase, two of the previously shown pictures were presented simultaneously next to each other and each within a grey frame. The picture pairs presented in the retrieval trial included the first and fourth and the second and fifth one of each list presented during the encoding phase, resulting in eight retrieval trials. Their arrangement on screen from left to right could match the sequential order in the list or not at random (50% chance) and the order was pseudorandomized. Participants were asked to indicate whether the arrangement from left to right of pictures in the pair matched the sequential order of the encoding phase and how confident they were about their answer (forced choice with four options). This resulted in eight retrieval trials per block. With 12 blocks, participants performed 96 retrievals with two pictures each, thus corresponding to 192 encoded stimuli for which subsequent memory retrieval was examined. The main behavioral outcome was percent correct answers.

### 2.3 EEG Recording

EEG was recorded during the sequence memory task using a 32-channel EEG setup (BrainProducts, Gilching, Germany) with active electrodes mounted to a cap according to the international 10-20 system, online high-pass filter of 0.016 Hz, a low-pass filter of 250 Hz, and 1000-Hz-sampling rate. Reference and ground electrode were placed on the nose tip and at FCz position, respectively. Four electrodes were mounted above and below the right eye and right and left outer canthi to record the electrooculogram (EOG). Impedances were kept below 5 kΩ with abrasive gel (Nuprep, Weaver and Company) for reference and EOG electrodes and Supervisc electrode gel (EasyCap, Herrsching, Germany) for all electrodes.

### 2.4 EEG Data Processing

EEG data processing was conducted using the Fieldtrip toolbox (Oostenveld et al., 2011) in Matlab (The MathWorks, Natick, MA). Due to technical failure, EEG data from one older participant was not stored. Channels containing no signal or substantial artefacts were rejected after visual inspection (1-4 channels for nine of the older adults, seven young adults with 1 rejected channel and one young adult with 6 rejected channels) and were interpolated. For three older participants the EOG channel above the right eye was replaced with Fp1 for vertical eye movements. An offline low-pass 50-Hz filter was applied. Then, the data was epoched into trials of 8-second-length (1.5 seconds before and 6.5 seconds after stimulus presentation during encoding). Epochs containing muscle activity were identified by visual inspection and removed. The remaining epoched data was down-sampled to 500 Hz, demeaned, and subsequently decomposed into 28 components with independent components analysis (ICA; Makeig et al., 1997). Components correlating (r ≥ .3 or r ≤ −.3) with the EOG signal converted to bipolar vertical and horizontal EOG were rejected (Hanna et al., 2014; Jung et al., 2000; Lucchese et al., 2017; Winkler et al., 2011). Remaining epochs containing voltage deflections above ± 150 µV were automatically rejected and only participants with a rejection rate below 25% were included in the following analyses (Lopez-Calderon and Luck, 2014; Luck, 2014), resulting in 21 older and 26 young adults (mean rejected trials (SD) young adults = 12.2 % (5.7), older adults = 12.7 % (6.5)).

Rejection rates did not differ between correct and incorrect trials in young (mean rejection rate correct trials (SD) = 12.2 % (6.0), incorrect (SD) = 10.8 % (9.7), *t*(25) = 0.70, *p* = .490) and older participants (mean rejection rate correct trials (SD) = 12.5 % (6.5), incorrect (SD) = 10.3 % (8.8), *t*(20) = 1.79, *p* = .089). Scalp current density was computed for the remaining data using the spherical spline method (Perrin et al., 1989). Then, time-frequency analysis was conducted for each epoch in a frequency range from 1 to 45 Hz in frequency bins of 0.5 Hz using a Morlet wavelet with varying number of cycles increasing from 3 to 10 (wavelet length *m* = 3; normalization factor *A* = σt−1/2 π−1/4) for time bins of 0.01 seconds (Garagnani et al., 2016; Roach and Mathalon, 2008; Schneider et al., 2008). Frequency spectra were averaged over correct and incorrect trials separately for each participant. Then, power spectra for each group and condition were decibel-baseline corrected with a baseline time window from 0.5 to 0.1 seconds pre-stimulus. Then, trials were separated into an early (0.1-2.9 sec) and late (3.1-5.9 sec) time window to temporally distinguish more perceptual processes during the beginning of a trial from more binding-related processes during the second phase of the trial (e.g., through the instructed strategy to associate each stimulus with the stimulus shown before) (Griffiths et al., 2021). To determine phase-amplitude coupling, the modulation index (MI; Tort et al., 2010) was calculated between theta (4-8 Hz) phase and gamma (30-45 Hz) amplitude.

### 2.5 Statistical Analysis

A two-sided alpha-level of 0.05 was applied for all tests. Behavioral data was analyzed using *R* software (R Core Team, 2019). Memory performance (percent correct answers) was compared between young and older participants using an independent sample *t*-test.

Statistical analyses of EEG data were performed using the Fieldtrip (Oostenveld et al., 2011) toolboxes in Matlab (The MathWorks, Natick, MA). We analyzed the interaction between age group and memory performance as illustrated in “How to test an interaction effect using cluster-based permutation tests?” on the Fieldtrip-Website (https://www.fieldtriptoolbox.org/faq/how_can_i_test_an_interaction_effect_using_cluster-based_permutation_tests/, last retrieval April 11, 2023). First, frequency spectra from incorrect trials were subtracted from correct trials for each group separately. Then, these group-wise difference scores were compared with an independent *t*-test. This comparison of differences between groups tested the interaction between memory performance and age group. Then, cluster-based permutation (cluster threshold = ± 1.96, 1000 permutations, minimum neighbourhood size = 2; Maris and Oostenveld, 2007) of *t*-values from each channel-frequency-timepoint combination was performed to determine the significance level of the interaction effect. As cluster-based permutation only tests the significance of cluster size and not their extent in time, frequencies or location (Groppe et al., 2011; Maris, 2011; Maris and Oostenveld, 2007; Sassenhagen and Draschkow, 2019), we performed cluster-based permutation separately for a-priori defined frequency bands [theta (4-8 Hz), alpha/beta (10-30 Hz), and gamma (30-45 Hz)] and time windows (early: 0.1-2.9 sec and late: 3.1-5.9 sec) to test our hypotheses regarding specific frequencies and time points. One young participant had only correct trials and was therefore not included in any analyses comparing correct and incorrect trials.

Additionally, similar to Heusser et al. (2016), a modelled MI with decreasing values for increasing picture positions was used as regressor to predict actual MIs (averaged over theta and gamma frequencies) for each trial, sensor and subject. Note that similar to other studies (Axmacher et al., 2010; Daume et al., 2017; Heusser et al., 2016; Reinhart and Nguyen, 2019; Tort et al., 2010), we calculated PAC on whole-trial data in all theta (4-8 Hz) and gamma (30-45 Hz) frequencies derived using Morlet wavelets (see details of frequency analysis above). As we were interested in the modulation of gamma amplitude by theta phase and not the magnitude of coupling per se, we did not determine sensors, trials or time points showing significant coupling first. Resulting *t*-values from the regression of predicted on actual MIs were included in a cluster-based permutation analysis separately for each age group, to determine electrodes showing the hypothesized pattern of MI. Here, picture position was randomly shuffled 1000 times for each participant, and the corresponding *t*-value for the modelled MI was calculated for each electrode. Resulting *t*-values were averaged over participants for each permutation, transformed into *z*-values and thresholded at *p* < .05. Maximum cluster size was determined for each permutation and used as threshold for the *t*-values from the measured MIs. To examine the pattern of coupling of gamma amplitude to theta phase, values from significant clusters were extracted for follow-up analyses and averaged over electrodes. To determine the preferred phase angle for each picture position depicted in the polar plots of ***Figure 4***, the main effect of coupling (coupling of gamma power to theta phase irrespective of picture position) was removed first. For this, we first calculated the normalized gamma power for each phase bin by dividing its mean gamma amplitude by the summed gamma amplitude over all bins. Then, we subtracted the mean normalized gamma power over all picture positions from the normalized gamma power of each picture position, separately for each phase bin. Next, individual phase bins for each picture position were determined by extracting the theta phase bin with the highest normalized gamma amplitude.

## Code accessibility

All analysis scripts are available from the corresponding author upon reasonable request.

## 3. Results

### 3.1 Memory performance

Older adults had significantly less correct answers than young adults (mean % correct older adults (SD) = 68.15 (6.83), young adults = 87.9 (7.22); *t*(45) = -9.57, *p* < .001).

### 3.2 EEG

#### 3.2.1 Theta and gamma power are differentially associated with memory performance in young and older adults

We found a significant interaction effect of age and memory performance on theta power during memory encoding. Cluster-based permutation revealed a significant difference between older and young adults regarding the difference between correct and incorrect trials (mean *t*-statistic within cluster = -2.54, *p*_corr_ = .01, cluster size = 406) in the early time window, suggesting that perceptual processes important for correct sequence encoding are affected by aging. The difference was most pronounced over left temporo-parietal electrodes and extended to frontal and central regions (see ***Figure 2A***). Specifically, post-hoc tests revealed that there was no significant difference between correct and incorrect trials in theta desynchronization in older adults. However, young adults showed significantly more desynchronization in the theta band during correct than incorrect trials (*p* = .002). There was no significant interaction in the theta band in the late time window (*p*_corr_ = .47). Testing for the interaction in the alpha/beta frequency band, no significant effects emerged for either time window (*p*’s_corr_ > .08). Frequency spectra showed no difference between correct and incorrect trials in alpha/beta power in older adults, while there was a tendency towards more alpha/beta power in correct than incorrect trials in young adults, especially in higher frequencies (see ***Figure 2B***). In the gamma band, a significant interaction effect was revealed in the late time window (mean *t*-statistic within cluster = 2.61, *p*_corr_ = .02, cluster size = 331), suggesting that also more binding-related processes important for correct sequence encoding are affected by aging. This effect was strongest over right frontal and central electrodes (see ***Figure 2C***). Post-hoc tests showed that there was no significant difference between correct and incorrect trials in older adults, while young adults had significantly more gamma power during correct compared to incorrect trials (*p* < .001). Additionally, young adults showed significantly less gamma power during incorrect trials than older adults (*p* = .004). There was no significant interaction effect in the gamma band in the early time window (*p*_corr_ = .55). In sum, we found differential effects of age on the subsequent memory effect in lower frequencies in the early time window (during presumably more perceptual processes) and in higher frequencies in the late time window (during presumably more memory-specific processes). In older adults, there were no significant differences between correct and incorrect trials in theta and gamma frequency and there was a tendency towards the opposite activation pattern compared to young adults. Together, these results indicate that the interplay between theta and gamma oscillations during sequence encoding is altered with aging.

**Figure 2.**
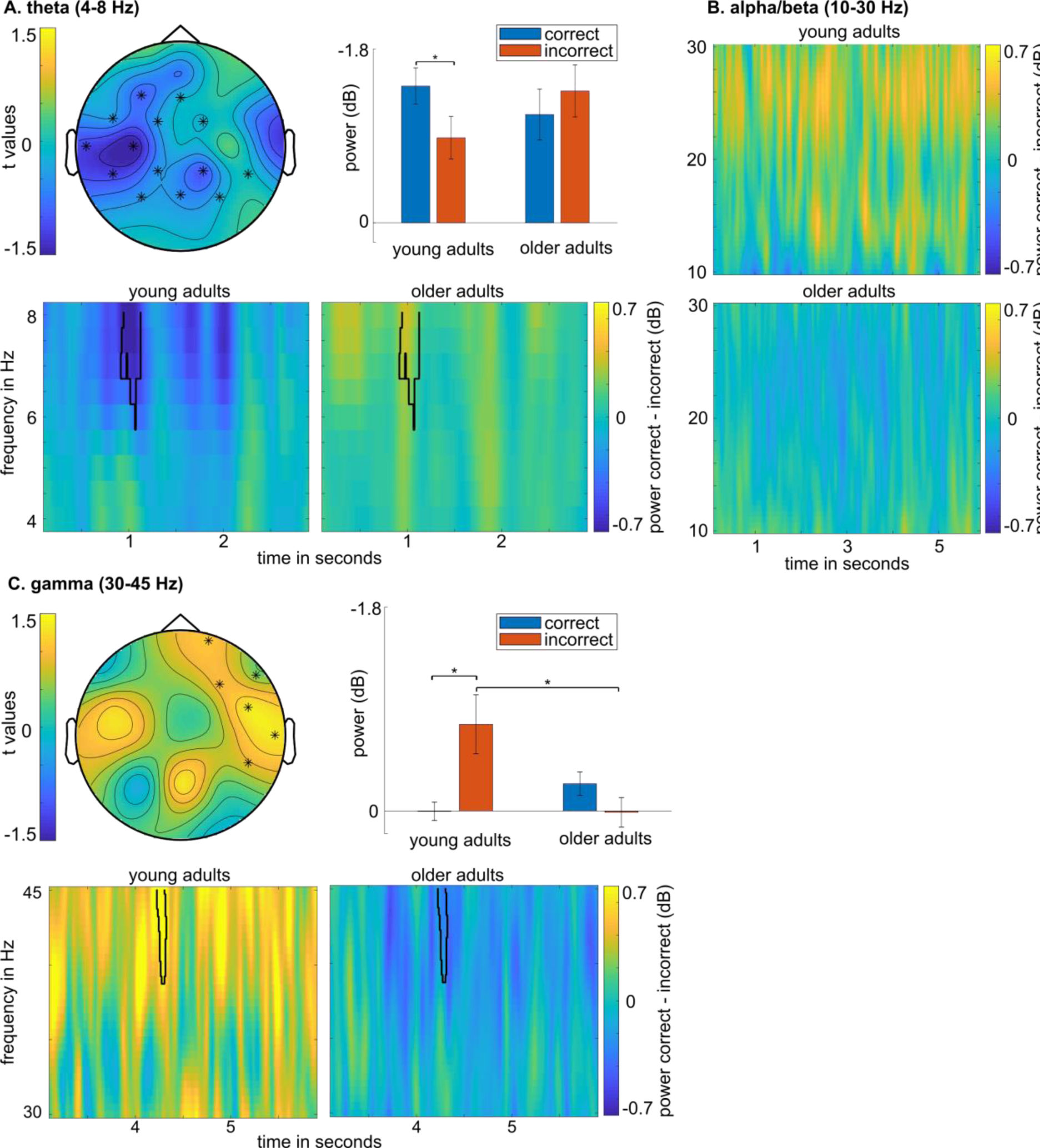
Panel A shows the significant interaction effect between age (young vs. old) and performance (correct vs. incorrect) in the early time window (0.1 – 2.9 sec) and theta (4 – 8 Hz) frequency band (*p*_corr_ = .01). The topography depicts the *t*-values of the comparison between correct-incorrect difference scores for older and young adults averaged over theta frequencies and timepoints 0.5 – 1.5 sec with stars indicating the significant cluster (*p*_corr_ < .05). *Notes.* Hz = Hertz; sec = seconds; dB = decibel.

The bar graph depicts average dB-corrected power values in the significant cluster for young and older adults for correct and incorrect trials each. Power spectra depict the baseline-corrected (dB-correction 0.5 to 0.1 sec pre-stimulus) difference between correct and incorrect trials for young (left) and older adults (right) with the cluster of the significant interaction effect outlined in black. Panel B shows the baseline-corrected (dB-correction 0.5 to 0.1 sec pre-stimulus) difference between correct and incorrect trials for young (top) and older adults (bottom) in alpha/beta (10-30 Hz) frequency. Panel C shows the significant interaction effect in the late time window (3.1 – 2.9 sec) and gamma (30 – 45 Hz) frequency band (*p*_corr_ = .02). The topography depicts the *t*-values of the comparison between correct-incorrect difference scores for older and young adults averaged over gamma frequencies and timepoints 4 – 4.5 sec with stars indicating the significant cluster (*p*_corr_ < .05). The bar graph depicts average dB-corrected power values in the significant cluster for young and older adults for correct and incorrect trials each. Power spectra show the difference between correct and incorrect trials for young (left) and older adults (right) with the significant cluster of their comparison outlined in black.

#### 3.2.2 Theta-gamma PAC codes sequential order of items in older adults

In older adults, the modelled MI predicted the measured coupling values according to picture position (*t(*20) > 0.44, *p*_corr_ = .05; see ***Figure 3A***). Visual inspection of the MI from this cluster per picture position revealed that the MI indeed slightly decreased with item position within a sequence (see ***Fig. 3B***), indicating that the later an item appears in a sequence, the wider gamma activity is spread over theta phases.

**Figure 3.**
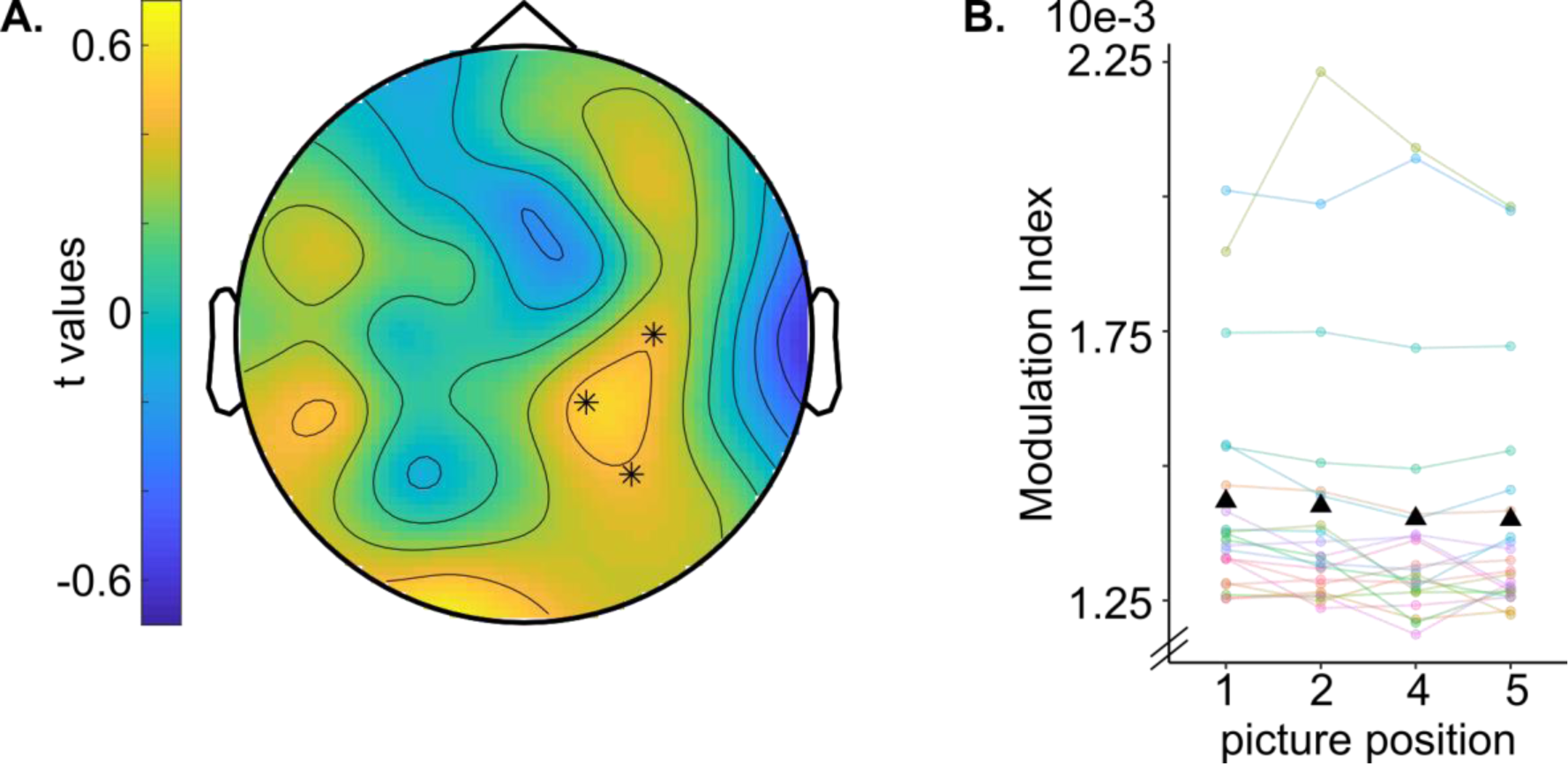
Panel A shows the topography of *t*-values resulting from regressing a modelled modulation index (MI) with decreasing values for increasing picture positions on measured MIs between gamma amplitude and theta phase during encoding. Stars indicate the significant cluster (*p*_corr_ ≤ .05). Panel B shows the MI between gamma (30-45 Hz) amplitude and theta (4-8 Hz) during the whole encoding trial (0.1-5.9 sec) averaged over electrodes in the significant cluster. Black triangles show the mean MI per picture position. Individual data points are depicted with colored dots and lines. Note that the MI for picture position 3 is not depicted because this picture was not included in the retrieval phase.

Furthermore, we inspected the distribution of gamma (30-45 Hz) amplitude over theta (4-8 Hz) phases, separately for later correctly and incorrectly remembered trials (see ***Figure 4***). For correctly remembered sequences, gamma amplitude during encoding of pictures presented early in a sequence was highest during early theta phases, while gamma amplitude during encoding of later pictures within a sequence was highest at later phases of theta. Crucially, for incorrectly remembered sequences, this modulation of gamma power by theta phase according to picture position was missing, indicating that the sequential nesting of gamma oscillations within theta cycles might be crucial for remembering the temporal order of items within a sequence. In young adults, no significant cluster emerged for the modelled MI representing picture position (*t*(25) < 0.44 *p*_corr_ ≥ 0.1), suggesting that there was no differential coupling of picture position between theta phase and gamma amplitude in the investigated frequency bands (see ***Supplementary Material*** for a figure of the results in young adults).

**Figure 4.**
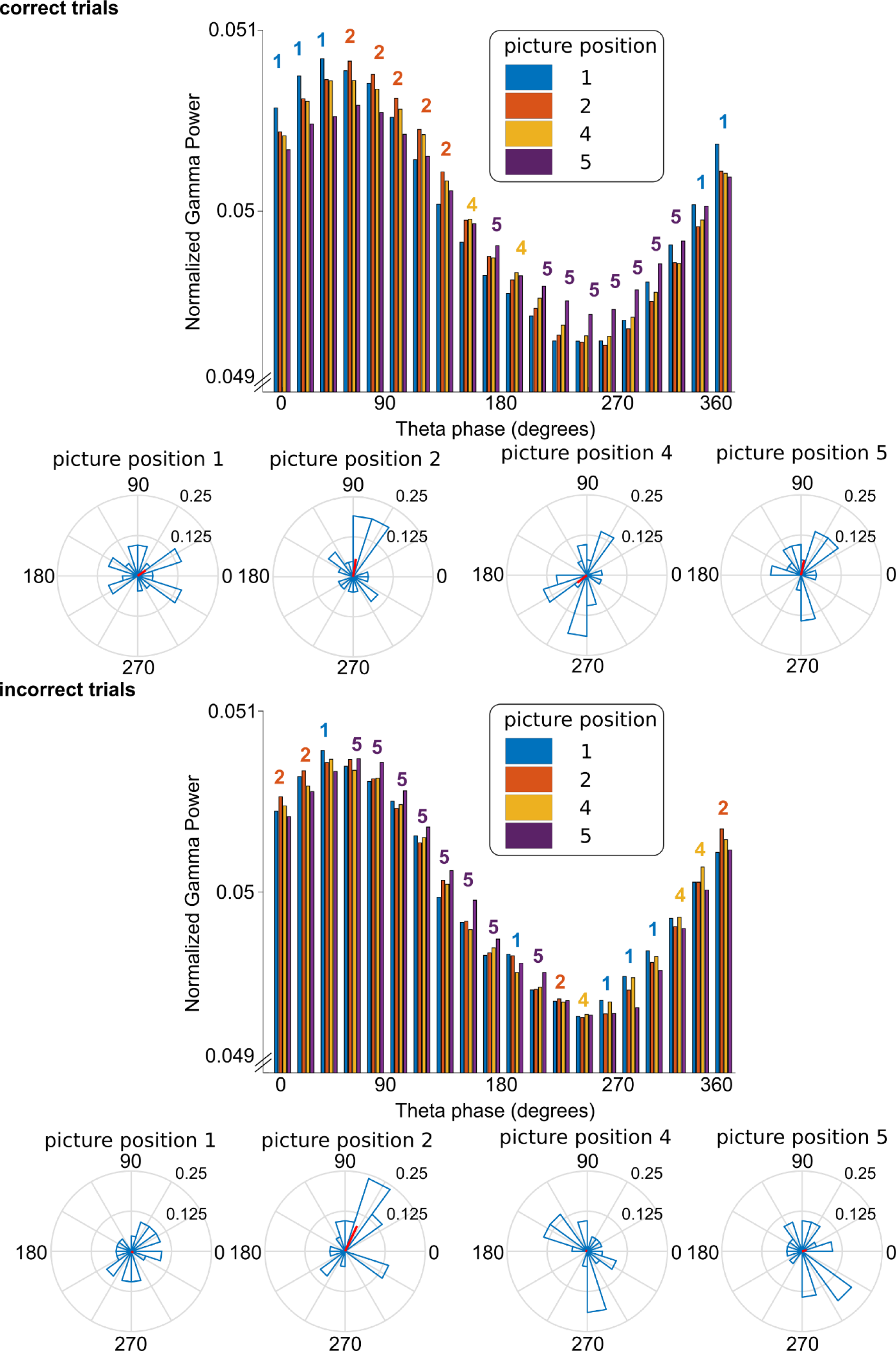
The bar graphs show the distribution of normalized gamma (30-45 Hz) amplitude (averaged over participants and electrodes in the significant cluster in Figure 3) over theta (4-8 Hz) phase during encoding per position of the picture within a sequence for later correctly (top) and incorrectly (bottom) remembered trials. The colored numbers above the bars indicate for which picture position gamma power was highest during each theta phase bin. For pictures presented early within a sequence, gamma amplitude was highest during early theta phases, while for pictures presented later in a sequence it was highest at later phases of theta. Crucially, for incorrectly remembered sequences (bottom), this modulation of gamma power by theta phase according to picture position was missing. The polar plots under the bar plots depict the distribution of preferred theta bins (phase bin with highest gamma power) for each picture position. For correctly remembered trials, the mean preferred phase increases with picture position within a sequence (top), while the preferred theta phase is less clear for incorrectly remembered trials (bottom).

## 4. Discussion

In this study, we aimed to investigate how oscillatory processes in different frequencies contribute to successful sequence encoding in older adults. We found that age group (older vs young adults) significantly affected EEG power in lower and higher frequencies during correct encoding of sequential stimuli. Specifically, in line with the Sync/Desync-model, a desynchronization in low frequencies (theta, 4-8 Hz) was beneficial for memory performance in young adults, followed by detrimental effects of desynchronization in gamma frequency (30-45 Hz) on memory performance. In older adults, even though this differential modulation of power was missing, we could show that correctly remembered sequences were still coded through modulation of gamma amplitude by theta phase.

The Sync/Desync model suggests that desynchronization in lower frequencies is important for correct encoding in young adults, because it allows for representation and processing of to-be-remembered stimuli through disinhibition of the cortex (Crespo-García et al., 2016; Hanslmayr et al., 2016; Hanslmayr et al., 2012). We found that this mechanism seems to be altered in older adults, with no significant difference in desynchronization between later correctly and incorrectly remembered trials in lower frequencies. Moreover, our results from young adults support the model, given that theta power during the encoding of subsequently correctly remembered trials was lower than during the encoding of subsequently incorrectly remembered trials. Although it has been suggested that decreases in low frequencies extend into alpha and beta bands, here we found a subsequent memory effect only in the theta band. Nevertheless, although we did not formally distinguish between a perceptual and a binding phase in the task, we found this interaction only in the first three seconds after stimulus onset. This is in line with a more perceptual/attentional mechanism that may underlie desynchronization in lower frequency bands (Griffiths et al., 2021; Hanslmayr et al., 2016; Hanslmayr et al., 2012).

In a recent review, it has been hypothesized that the subsequent memory contrast that we applied here captures not only memory-specific, but also more general perceptual and attentional processes that are important for correct encoding of to-be-remembered stimuli (Herweg et al., 2020). However, not only lower frequencies commonly associated with such perceptual/attentional processes were differentially affected by age and memory performance, but also higher frequencies, i.e., gamma power, showed an age group-specific pattern. Gamma spikes are thought to enable perceptual binding processes and represent processed items in the cortex as input to the hippocampus (Lisman and Jensen, 2013; Nyhus and Curran, 2010). Accordingly, increased gamma power has been associated with memory performance (Friese et al., 2013; Osipova et al., 2006; Sederberg et al., 2006). However, in older adults there was no difference in gamma power between later correctly and incorrectly remembered items, suggesting that the to-be-remembered items might not be adequately represented in the cortex by gamma oscillations. In young adults, desynchronization in the gamma frequency band was detrimental for encoding of sequential stimuli in young adults. In line with the Sync/Desync model, the significant effects in gamma power occurred during a later phase of encoding, where participants were presumably engaged in more memory-specific processes like mnemonic binding (Griffiths et al., 2021).

Thus, our results in young adults are in line with models suggesting a decrease in lower frequencies that enable efficient processing of stimuli through synchronization in higher frequencies, and extend these models for sequence memory. Moreover, our data suggests that these complementary desynchronization and synchronization processes in relation to correct encoding of sequential stimuli is not fully preserved in older adults. Namely, the missing power difference in theta and gamma frequency bands between later correctly and incorrectly remembered trials in older adults might point towards neural dedifferentiation, i.e. less distinct neural representations of to-be-remembered items (Abdulrahman et al., 2017; Koen et al., 2019; Li et al., 2001), which may explain some age-related variance in episodic memory (Koen et al., 2019). This finding is in line with previous results showing more similar frequency-representations of items in older than in young adults which correlated with associative memory, while dissimilarity in neural representations was advantageous for memory performance in young adults (Sommer et al., 2019).

Additionally, for the task used in the present study, not only encoding of the stimulus per se is essential, but more importantly the encoding of its position within a sequence. Therefore - in addition to power changes in single frequency bands – we looked at the nesting of gamma oscillations in theta phase to disentangle the oscillatory correlates of sequence encoding. For older adults, a significant cluster showing the hypothesized pattern of MI emerged in right central and temporal electrodes. Visual inspection of the distribution of gamma power over theta phase in older adults showed that gamma power was highest in an earlier theta phase for pictures that were presented early in a sequence, while it was highest in later theta phases for pictures later in a sequence. Thus, our results replicate the coding mechanism for temporal order in a sequence of Heusser et al. (2016) in healthy older adults, suggesting a similar process for correct encoding of sequences than in young adults, albeit in a lower gamma frequency range. A general slowing of neural oscillations with age has been found (Courtney and Hinault, 2021), possibly explaining why we found this pattern of theta-gamma PAC in the low gamma frequency band (30-45 Hz) in older adults. In young adults, the modulation of theta-gamma PAC by position within a sequence has been found in a higher gamma frequency (70-100 Hz) (Heusser et al., 2016) that we could not analyze due to high-frequency artifacts in our data. Additionally, we adapted the sequence memory paradigm of (Heusser et al., 2016) specifically for older adults, possibly further explaining the discrepancy in our findings in young adults. Crucially, older adults presented significantly more incorrect trials than young adults, in which the distribution of gamma power over theta phase was not according to picture position. Therefore, this theta-gamma coding of sequences might lose precision in healthy aging, providing a possible explanation for age-related decline in sequence memory. Although decreased theta-gamma PAC in older compared to young adults has been found for associative and working memory (Karlsson et al., 2022; Reinhart and Nguyen, 2019), this is the first time, to our knowledge, that it has been investigated specifically for the coding of sequences.

### 4.1 Conclusion

Together, our results provide a putative explanation for age-related decline in sequence memory. Our results are in line with models suggesting decreased activity in low frequency bands, increase of activity in high frequency bands, and its nesting to lower frequencies, as key mechanisms to enable efficient information processing and memory-specific binding of items for successful encoding of stimuli into episodic memory. Here, we extend these findings to sequence memory in young and older adults. In older adults, both desynchronization in lower and synchronization in higher frequencies are disturbed, potentially reflecting a mechanism underlying deficient sequence memory. Crucially, while the coding of sequences through the sequential nesting of gamma oscillations in theta waves seems to be generally intact, it might be less precise in older adults, leading to many incorrectly remembered sequences. Future studies could directly test these proposed mechanisms of age-related sequence memory decline and possibilities to improve sequence memory in older adults using transcranial alternating current stimulation (tACS). Additionally, studies in individuals with memory decline, e. g., patients with mild cognitive impairment, may elucidate the clinical relevance of this process in memory decline.

## Author contributions

NME: data curation, formal analysis, investigation, visualization, writing – original draft, writing – review & editing. AF: conceptualization, funding acquisition, methodology, resources, supervision, writing – review & editing. S-CL: methodology, writing – review & editing. GL: conceptualization, formal analysis, methodology, supervision, writing – review & editing. DA: conceptualization, funding acquisition, methodology, project administration, resources, supervision, writing - review & editing.

## Acknowledgements

This work was funded by the Deutsche Forschungsgemeinschaft (DFG, German Research Foundation) – 426477764 (to DA and AF) and by a “Gerhard Domagk” research grant awarded by the University of Greifswald Medical School (to GL). We thank Robert Malinowski for programming the task and technical support with the EEG equipment, all our involved student assistants for their help with data acquisition and all participants taking part in the study.

## Websites

Fieldtrip Toolbox. Retrieved April 11 2023 from https://www.fieldtriptoolbox.org/faq/how_can_i_test_an_interaction_effect_using_cluster-based_permutation_tests/

## Notes

### Competing Interest Statement

The authors have declared no competing interest.

