## Supplementary Material for "Brain oscillatory processes related to sequence memory in healthy older adults"

***Supplementary Table.*** Neuropsychological assessment of 21 healthy older adults.

| | Mean (SD) |
|---|---|
| Geriatric depression scale | 0.90 (0.94) |
| Semantic fluency (number of animals) | 26.19 (6.31) |
| Boston Naming Test (max. 15) | 14.57 (0.68) |
| Mini-Mental State Examination | 29.33 (0.80) |
| Word list learning | |
| Total (max. 30) | 23.33 (2.96) |
| Trial 1 (max. 10) | 6.19 (1.25) |
| Trial 2 (max. 10) | 8.10 (1.26) |
| Trial 3 (max. 10) | 9.05 (0.80) |
| Word list retrieval | 8.57 (1.33) |
| Word list intrusions | 0.19 (0.40) |
| Figure copying (max. 11) | 10.95 (0.22) |
| Figure retrieval (max. 11) | 10.67 (0.97) |
| Phonematic fluency (number of s-words) | 16.43 (4.09) |
| Trail-making test | |
| Part A (sec) | 40.20 (11.42) |
| Part B (sec) | 81.55 (14.34) |
| Digit-span | |
| Forward | 9.05 (2.73) |
| Backward | 7.19 (1.86) |
| Identical pictures | |
| Number correct responses | 21.24 (2.36) |

| | |
|---|---|
| RT (ms) | 3363.24 (1008.24) |
| Spot-a-word | |
| Number correct responses | 24.14 (5.44) |
| RT (ms) | 6089.87 (3673.44) |

*Note*. SD = standard deviation. Sec = seconds. Ms = milliseconds.

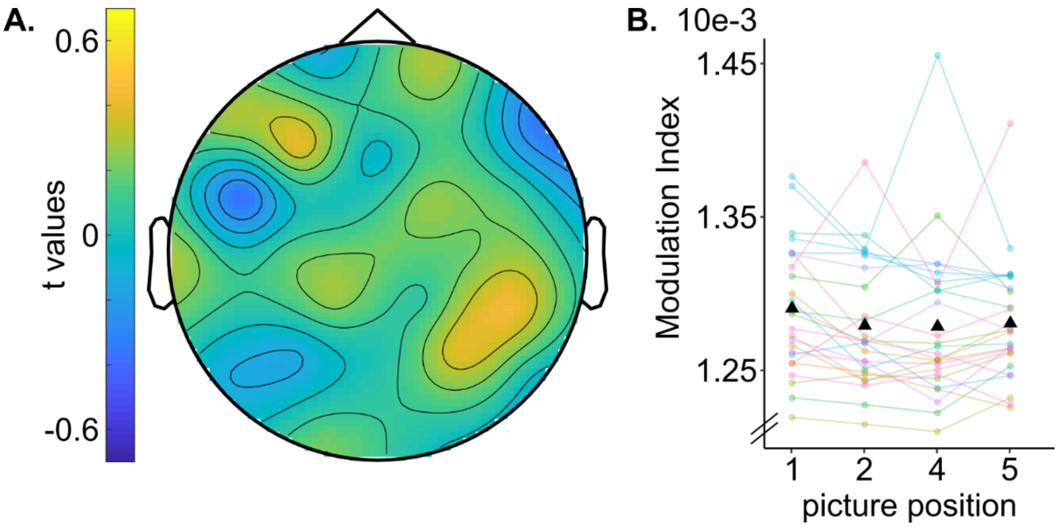


***Supplementary Figure.*** Panel A shows the topography of *t*-values resulting from regressing a modelled modulation index (MI) with decreasing values for increasing picture positions on measured MIs between gamma amplitude and theta phase during encoding. There were no significant clusters ($t(25) < 0.44$ $p_{corr} \geq 0.1$). Panel B shows the MI between gamma (30-45 Hz) amplitude and theta (4-8 Hz) during the whole encoding trial (0.1-5.9 sec) averaged over all electrodes. Black triangles show the mean MI per picture position. Individual data points are depicted with coloured dots and lines. Note that the MI for picture position 3 is not depicted because this picture was not included in the retrieval phase.
